# Structural basis of SARS-Cov-2 spike recognition by engineered synthetic multivalent VHH antibodies

**DOI:** 10.1101/2024.10.02.616254

**Authors:** Ana G. Lujan Hernandez, Zane T. Laughlin, Anamika Patel, Tom Z. Yuan, Rebecca L. Nugent, Fumiko Axelrod, Eric A. Ortlund, Aaron K. Sato

**Author notes:** AGLH and ZTL contributed equally to this work. To whom correspondence should be addressed: Aaron Sato, Eric Ortlund.

## Abstract

High-throughput technologies such as next-generation sequencing (NGS), microarray-based gene synthesis, and phage display have empowered the discovery and engineering of precisely defined, synthetic antibodies with high avidity and drug-like features. Here, we describe a scalable process for engineering homo- and hetero-hexavalent variable domains of camelid heavy-chain (VHH)-Fc antibodies against the severe acute respiratory coronavirus 2 (SARS-CoV-2) spike (S) protein. Overall, we demonstrate that VHH trimerization is an effective and modular approach for increasing the affinity of anti-S1 VHH-Fc antibodies for the highly mutated S proteins of SARS-CoV-2 variants. We show that one specific nanobody (named TB201-1) binds spike trimer protein at the interface of two neighboring RBDs, recognizing one distinct epitope on one RBD but making a set of secondary interactions with the neighboring RBD. From this structure, we determine the epitope-paratope residues responsible for spike-nanobody interaction and how mutations found in the SARS-CoV-2 variants contribute to oblate TB201-1 binding. This approach could be leveraged to improve existing antibody-based diagnostics and therapeutics targeting SARS-CoV-2 as the virus evolves.

## Introduction

Antibodies must bind their targets with high avidity to affect therapeutic or diagnostic outcomes. This parameter is specified by the complementarity-determining regions (CDRs) of an antibody’s paratope, or antigen-binding site. The mammalian immune system can produce antibodies with micro- to nanomolar affinity through *in vivo* affinity maturation via somatic hypermutation, but higher affinities (<100 pM) are precluded by the physiological constraints of B cell activation.^1^ Methods for *in vitro* affinity maturation have surpassed this affinity ceiling, enabling the production of antibodies with low picomolar and femtomolar affinities.^2^ Although computational approaches have been proven successful in this regard, they are less scalable than *in vitro* display technologies, which can be applied iteratively to select for higher and higher affinity binders from large antibody libraries. *In vitro* display technologies also offer immediate access to the antibody fragment genes, which can subsequently be multimerized to generate high-avidity antibody fusions.

Among the available antibody formats, single-domain antibodies such as variable domain of camelid heavy-chain (VHH) antibodies (also known as nanobodies) are one of the most engineerable. Lacking a light chain, VHH antibodies are very small at ∼12-15 kDa — half the size of single-chain variable fragment (scFv) antibodies, a third the size of fragment antigen-binding (Fab) antibodies, and less than a tenth the size of immunoglobulin Gs (IgGs). Their small size confers several benefits including targeting of occluded and non-immunogenic epitopes normally inaccessible to other antibody formats^32^. Moreover, VHHs are readily expressible on bacteriophage M13 and in *E. coli*, permitting affinity maturation by phage display and high-throughput screening of engineered multivalent VHHs. Other antibody formats, such as scFv and Fab, can also be expressed in these microorganisms, but are more difficult to express in multivalent formats.^3^ Even when VHHs are joined in tandem, the resulting multivalent constructs retain their small size and therefore, have advantages over larger antibody formats. These advantages have encouraged many drug developers to prioritize VHHs when developing diagnostic and therapeutic antibodies against severe acute respiratory coronavirus 2 (SARS-CoV-2), the etiological agent of the coronavirus disease 2019 (COVID-19) pandemic.^4^ In the first 18 months of the pandemic, scores of anti-SARS-CoV-2 single-domain antibodies were identified by phage display, including many with neutralizing activity and sub-nanomolar affinity for the spike (S) protein.^5–13^ Multivalent SARS-CoV-2 antibodies have been engineered more recently.^8,9,14–19^

VHH antibodies can be sourced from immune, naïve, or synthetic antibody libraries. Immune and naïve VHH libraries are generated by amplifying the VHH genes of immunized and unimmunized camelids, respectively, using real-time PCR (RT-PCR). The procedure, however, necessitates the use of multiple animals to generate a highly diverse library, as each individual camelid will have a unique antigenic response.^20^ Moreover, the repertoires from which immune and naïve libraries are sourced are redundant and biased, limiting the diversity that can be attained in such libraries.^21^ Synthetic libraries, which use synthetic oligonucleotides to introduce sequence degeneracy within the CDRs, overcome this limit.^22^ Advances in oligonucleotide and gene synthesis technologies have made the fabrication of massive synthetic libraries straightforward with high sequence diversity (>10^10^ clones), accuracy (1 error per 2 kb), and productivity (generally greater than 80% properly encoded sequences). We have generated a series of fully synthetic antibody libraries using a commercial DNA synthesis platform capable of synthesizing 1 million 300-mers on a single silicon chip (Twist Bioscience, patent WO 2015021080). In our approach, synthetic oligonucleotides encoding CDRs 1, 2, and 3 are synthesized and assembled into full-length VHH genes, which are then cloned into a vector for phage display. Recently, we reported the discovery of several high-affinity VHH-Fc antibodies against the SARS-CoV-2 S1 protein subunit from these libraries.^23,24^

Unlike methods for the construction of synthetic antibody libraries, methods for screening multivalent constructs are comparatively low throughput. Rational cocktail screening methods for monovalent binders have been leveraged to engineer multivalent constructs.^8,9^ While generally effective for lower-order multivalent constructs, these approaches are biased and difficult to apply to higher-order multivalent constructs where the number of possible permutations increases exponentially. In this paper, we developed an unbiased, high-throughput strategy for engineering synthetic multivalent VHH antibodies at scale. Beginning with 13 anti-S1 VHH leads,^24^ we constructed multivalent libraries of tandem-assembled VHHs, screened them by phage display, and sequenced the enriched libraries to determine the positional frequencies of each individual VHH fragment within the trimeric assemblies. Using this pipeline, we engineered mono- and bispecific multivalent VHH-Fc constructs with higher affinity for S1 than their monovalent counterparts and identified one tight VHH binder (TB201-1) that bound spike trimer protein with a K_D_ of 0.082 nM. To further define its epitope on spike protein, we determined the structure of TB201-1 in complex with spike trimer protein by single particle cryogenic electron microscopy (cryoEM). To our surprise, VHH TB201-1 binds the receptor binding domain (RBD) both in “up” and “down” conformation at the interface in spike trimer even though it was raised against monomeric spike S1. Three distinct binding modes of VHH to spike trimer were determined in cryoEM data, with a primary epitope located in the RBD-1 epitope class^33^. This interaction surface was present despite the conformation of S1 within the trimer and was common in all binding modes, suggesting it is the primary epitope. However, within the binding mode-2, where a VHH antibody binds to the spike RBD in the down conformation, the antibody makes a secondary set of interactions with the neighboring RBD at the RBD-7 epitope surface. These interactions are not common to all classes and are thus thought to be secondary. This structural knowledge can be used to better understand the mechanisms of resistance to this antibody (and related antibodies) by SARS-CoV-2 variants, particularly the Omicron sub-variants.

## Results

### Generating high affinity SARS-CoV2 specific multimeric VHH cassettes

Building on our neutralizing, high-affinity anti-S1 VHH-Fc antibodies from phage-display panning using our in-house synthetic VHH libraries,^24^ we sought to further improve their affinity by trimerizing the VHH domain. These leads were identified from two phage libraries: TB201, a humanized llama nanobody library with shuffled, llama-based CDR diversity; and TB202, a humanized llama nanobody library constructed using natural llama CDR1/2 sequences and human CDR3s identified from human naïve and memory B cells.

We initially constructed homo-hexavalent VHH-Fc antibodies for each lead by ligating three identical VHH cassettes together and inserting the resulting construct into an Fc acceptor vector (**Figure 1a**) which then dimerizes to become hexavalent. We separated each VHH domain with a (GGGGS)_2_AS linker. The trimeric nature of the constructs along with the repetitive and GC-rich nature of the linkers make the synthesis of full-length DNA constructs challenging. In particular, the presence of multiple long, high-repeat regions with high melting temperatures leads to high failure rates during gene synthesis.^25^ To overcome this challenge, we flanked VHH genes with Type IIS restriction sites, which recognize asymmetric DNA sequences and cleave outside of their recognition sequence, to enable Golden Gate assembly of three VHH fragments in tandem. Then we used homologous overhangs to enable the insertion of these trimeric constructs into an Fc-fusion acceptor vector, as illustrated in **Figure 1a**. Hexavalent VHH-Fc fusions were abbreviated according to the terminal identification number (e.g., “76” for TB202-76) and position of each VHH sequence in the tandem assembly (primary, secondary, or tertiary). For example, a homo-hexavalent construct containing three copies of the TB201-1 VHH gene is referred to as 1-1-1, whereas a hetero-hexavalent construct containing the TB202-76 VHH gene in the primary position and the TB201-1 VHH gene in the secondary and tertiary positions is referred to as 76-1-1.

**Figure 1.**
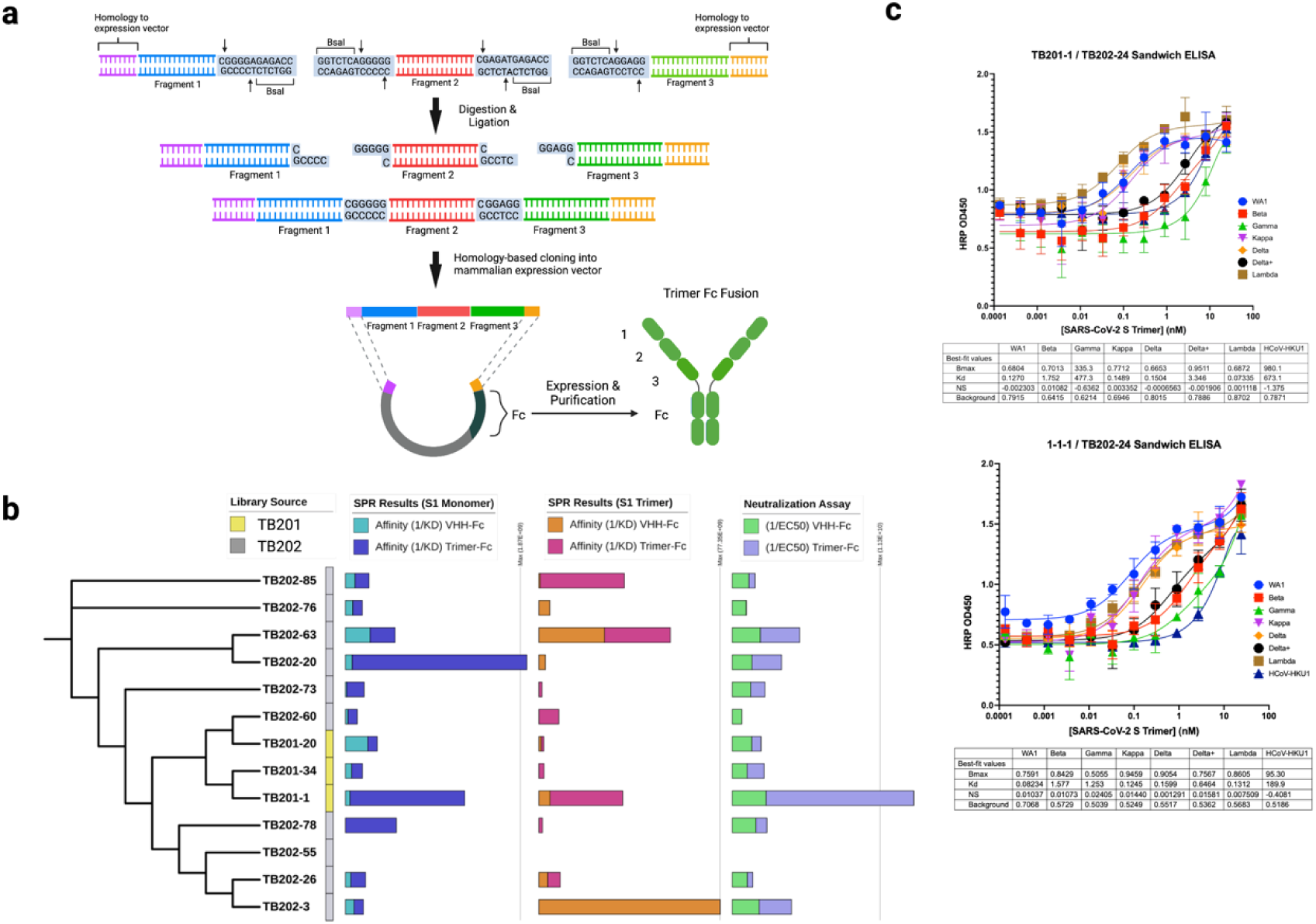
Screening homohexavalent anti-S1 VHH-Fc antibodies. (a) Homo-hexavalent anti-S1 VHH-Fc assembly strategy. (b) Comparative performance of homo-hexavalent VHH-Fc antibodies ([VHH]_3_-Fc) against their parental VHH-Fc counterparts in monomer S1 and S Trimer binding (SPR) and in S1-ACE2 competition assays. (c) Comparative performance of 1-1-1 and TB201-1 in sandwich ELISAs detecting the S trimer of the WA1, Beta, Gamma, Kappa, Delta, Delta+, and Lambda SARS-CoV-2 variants. Dissociation constants are presented in Table 1.

**Table 1.** Summary of dissociation constants (K^D^) between antibodies and SARS-CoV-2 S trimer variants, as measured by sandwich ELISA. ELISA experiments were performed in quadruplicate. Units are nanomolar (nM).

| Immobilized Antibody | WA1 | Beta | Gamma | Kappa | Delta | Delta+ | Lambda | HCoV-HKU1 |
| --- | --- | --- | --- | --- | --- | --- | --- | --- |
| 1-1-1 | 0.082 | 1.577 | 1.253 | 0.125 | 0.160 | 0.646 | 0.131 | 189.9 |
| TB201-1 | 0.127 | 1.752 | 477.3 | 0.149 | 0.150 | 3.346 | 0.073 | 673.1 |
| Fold Improvement | 1.54x | 1.11x | 380.93x | 1.2x | 0.94x | 5.18x | 0.56x | 3.54x |

### Affinity Studies

We initially compared homo-hexavalent VHH-Fc antibodies to their homo-bivalent VHH-Fc counterparts using surface plasmon resonance and a flow cytometry S1-angiotensin-converting enzyme 2 (ACE2) competition assay (see Methods). Overall, trivalent VHH sequences conferred higher binding affinities to the S1 monomer counterparts (TB202-20, TB201-1, TB202-60) and spike trimer (TB201-1, TB202-85), as well as greater competition with the ACE2 receptor in the S1-ACE2 competition assay (TB201-1) (**Figure 1b**). This result is presumably mediated by an avidity effect in a concerted binding to the three binding sites on the spike trimer. Whereas trivalency only improved one or two of these measures for most candidates, trivalent TB201-1 (1-1-1) resulted in marked improvements across all three measures. Not all antibody-S1 combinations produced SPR data (e.g., TB202-55).

To further characterize 1-1-1, we evaluated its ability to detect a wide variety of SARS-CoV-2 S variants when paired with TB202-24 in a sandwich ELISA. In this assay, immobilized 1-1-1 or TB201-1 served as a capture antibody; TB202-24 was then used as a detector antibody because it binds a unique S1 epitope not targeted by any other antibody described in this paper.^24^ After blocking, serial dilutions of various SARS-CoV-2 S trimers were detected using biotinylated TB202-24 and streptavidin-HRP (**Figure 1c**). S trimers from the ancestral WA1/2020 strain and the Beta, Gamma, Kappa, Delta, Delta+, and Lambda variants were assayed. The S trimer of human coronavirus HKU1 (hCoV-HKU1) was used as a negative control.

1-1-1 detected all S variant trimers except the Delta and Lambda S trimers with a higher apparent binding affinity than TB201-1 (**Figure 1c**, **Table 1**). The lack of improvement observed with the Delta and Lambda variants appeared to be a consequence of TB201-1’s existing high affinity for these S variants (0.15 and 0.073 nM, respectively). The most dramatic improvements in apparent binding affinity were observed with the Gamma and Delta+ S trimers, presumably because these variants were relatively weakly bound by TB201-1. The relatively weak association between TB201-1 and the Delta+, Beta, and Gamma S variants may be explained partly by the fact that all three of these variants possess a mutation at residue K417, which we previously identified as a key residue in TB201-1’s S1 binding epitope.^24^ However, it remains unclear why trivalent TB201-1 improved the detection of the Gamma and Delta+ S trimers but not the Beta S trimer. It is possible that a conformational change is required to enhance affinity to Gamma and Delta, which does not occur for the Beta S trimer.

Next we generated a phage library using our homotrimeric assembly strategy to interrogate the 2,197 homo- and heterotrimeric permutations that can be assembled from our initial 13 VHH leads (**Figure 2a**). The resulting phage library was panned for two rounds at a low antigen concentration to enrich for high-affinity binders. Subsequent long-read NGS revealed TB201-1 as the most prevalent VHH fragment at the secondary position within the linear tandem assembly after enrichment; the most frequent VHH fragments at primary and tertiary positions were TB201-1 and TB202-76 (**Figure 2b**). Initially we planned to select heterotrimeric constructs for further assessment based on enrichment per trimer library position (primary, secondary and tertiary), however the sequencing results revealed 1-1-1 homotrimer as a clear lead. Moreover, although there were other enriched clones per position, e.g. TB202-63 at the primary position and TB202-76 at the tertiary position, their enrichment was not as striking as that for TB201-1. Since TB202-76 at the primary position was the only clone to show preferential enrichment over TB201-1, we cloned the 76-1-1 bispecific hexavalent construct into a mammalian expression vector for further characterization.

**Figure 2.**
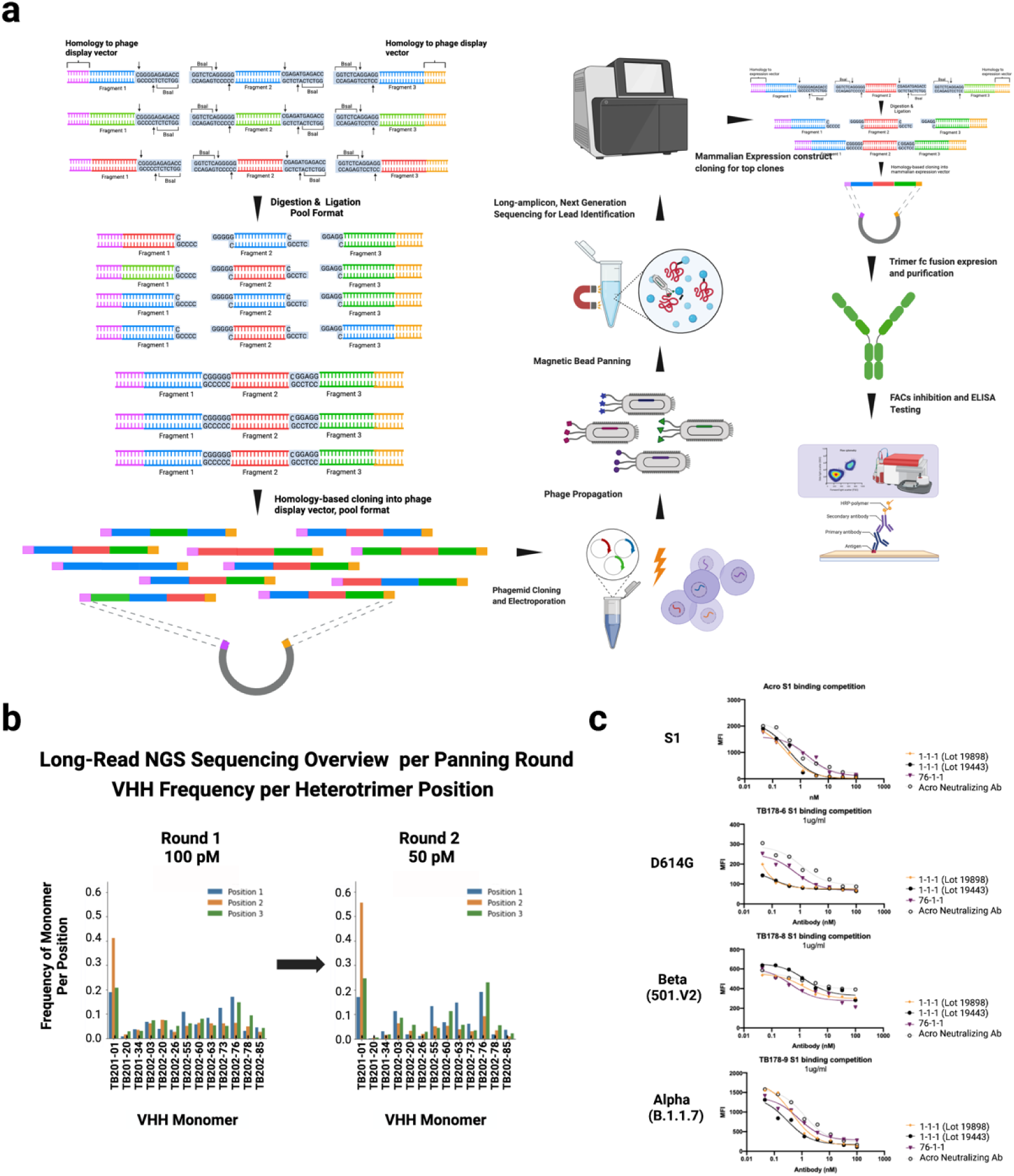
Screening heterohexavalent anti-S1 VHH-Fc antibodies. (a) Heterotrimeric VHH assembly and screening strategy. (b) Frequency of VHH sequences at each position (primary, secondary, and tertiary) within the tandem assembly, as determined by long-read NGS. (c) Comparative performance of 76-1-1 and two independently generated preparations of 1-1-1 in a S1-ACE2 competition assay incorporating S1 mutants from the D614G, Alpha, and Beta SARS-CoV-2 variants.

We initially used SPR to determine the apparent affinities between 1-1-1, 76-1-1, TB201-1, and TB202-76 and the S trimer of the ancestral SARS-CoV-2 strain and those of SARS-CoV-2 Alpha, Beta, C.1.2, Delta, Delta+, Lambda, Mu, and Omicron (**Table 2**). We also assayed affinities to the ancestral S1 monomer and the mutated S1 monomers of SARS-CoV-2 Delta and Omicron (**Table 3**). Interestingly, TB202-76 was the only construct that bound the Omicron S trimer (**Table 2**), and none of the constructs bound the Omicron S1 monomer (**Table 3**). As for the other S trimers, 1-1-1 demonstrated the highest apparent affinities (for all variants), followed by TB201-1, then 76-1-1, and finally TB202-76 (**Table 2**). The affinity improvement observed with the trivalent TB201-1 was modest, reflecting the already high (picomolar) affinity of TB201-1 for these SARS-CoV-2 S trimers. By contrast, adding two TB201-1 domains to TB202-76 (i.e., 76-1-1) resulted in a much greater affinity improvement for all S trimers except Omicron.

**Table 2.** Summary of apparent binding affinities between antibodies and SARS-CoV-2 S trimer variants, as measured by SPR. SPR experiments were performed in duplicate. Units are in picomolar (pM). n.b. = no binding observed.

| Antibody | SARS-CoV-2 S Trimer | SARS-CoV-2 S Trimer Alpha | SARS-CoV-2 S Trimer Beta | SARS-CoV-2 S Trimer C.1.2 | SARS-CoV-2 S Trimer Delta | SARS-CoV-2 S Trimer Delta+ | SARS-CoV-2 S Trimer Lambda | SARS-CoV-2 S Trimer Mu | SARS-CoV-2 S Trimer Omicron |
| --- | --- | --- | --- | --- | --- | --- | --- | --- | --- |
| TB201-1 | 13.52 ± 0.06 | 12.68 ± 0.13 | 507.13 ± 12.13 | 162.76 ± 9.06 | 19.65 ± 6.14 | 208.94 ± 2.3 | 12.64 ± 0.08 | 137.3 ± 8.85 | n.b. |
| TB202-76 | 7070 ± 573 | 5511 ± 129 | 21004 ± 2130 | 17214 ± 2281 | 13127 ± 722 | 26136 ± 3461 | 8354 ± 102 | 18683 ± 1932 | 25902 ± 7306 |
| 1-1-1 | 11.30 ± 0.58 | 6.93 ± 0.22 | 281.1 ± 4.67 | 95.20 ± 4.25 | 31.79 ± 2.19 | 73.56 ± 2.67 | 9.79 ± 2.09 | 93.89 ± 0.59 | n.b. |
| 76-1-1 | 84.63 ± 1.00 | 61.13 ± 1.16 | 1378.93 ± 70.52 | 212.63 ± 28.79 | 108.59 ± 0.53 | 233.32 ± 13.12 | 90.49 ± 10.86 | 186.07 ± 5.52 | n.b. |

**Table 3.**
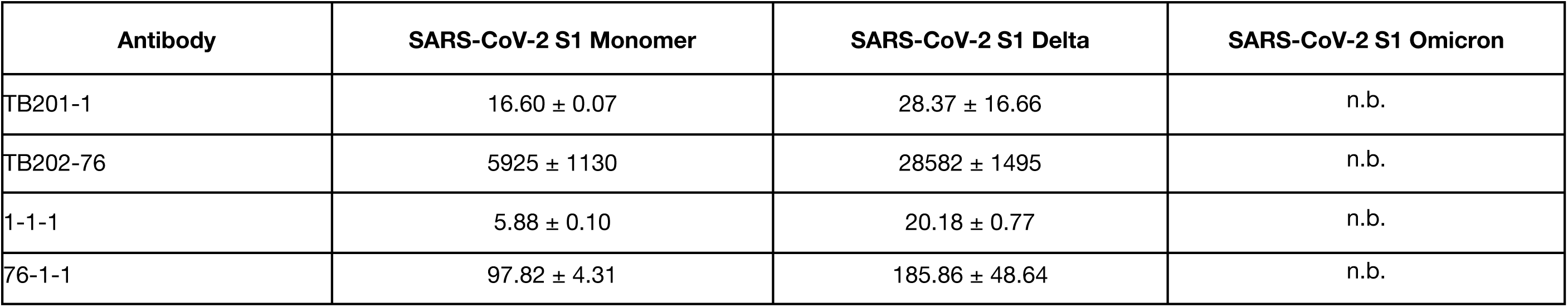
Summary of apparent binding affinities between antibodies and SARS-CoV-2 Ancestral, Delta, and Omicron S1 monomers, as measured by SPR. SPR experiments were performed in duplicate. Units are in picomolar (pM). n.b. = no binding observed.

We finally evaluated the ability of our lead monospecific (1-1-1) and bispecific (76-1-1) hexavalent VHH-Fc fusions to compete with cellular ACE2 for binding to mutant S1 proteins, including the D614G mutant and those of SARS-CoV-2 Alpha and Beta, in our flow cytometry assay. We also tested SAD-S35, a commercial anti-S antibody developed by ACROBiosystems, which behaved like TB201-1 in the binding, competition, and neutralization assays of our previous study.^26^ Whereas 1-1-1 exhibited greater potency against the wild-type, D614G, and Alpha S1 proteins, 76-1-1 more potently blocked the interaction between ACE2 and the S1 protein of the Beta variant (**Figure 2c**). Compared to SAD-S35, 1-1-1 demonstrated more potent competition with the ancestral S protein and the D614G and Alpha S1 variants, and 76-1-1 demonstrated more potent competition with the D614G, Alpha, and Beta S1 variants. Of the tested S1 proteins, the Beta variant is the only one that possesses a K417 mutation; thus, these data confirm the reduced performance of TB201-1 against S proteins containing a K417 mutation while buttressing bispecificity as a strategy for achieving broad-spectrum targeting of viral proteins.

### Structural Studies

We determined the structure of the TB201-1 (1-1-1) VHH in complex with the SARS-CoV-2 (Wuhan) spike protein trimer via single-particle cryoEM, revealing three distinct binding modes whereby the conformation of the spike trimer varied (**Figure 3-5, Supplementary Figure 1**). Across all three binding modes, the VHH was consistently found to recognize the epitope on the top RBD pocket, aligning with the RBD-1 epitope class as described earlier^33^. In binding mode 1, the VHH binds the RBD in the “up” conformation recognizing a single RBD surface (**Figure 3a, 5a, S4a**). Residues F486 and Y489 are the primary target residues in a binding site located on the RBD loop comprising residues 485-489 (**Figure 3b, 5b**). Spike F486 is recognized by VHH residues F37 (located on a beta-strand between CDR-1 and -2) and F105 (located on CDR-3) **(Figure 3c, 5c**). Spike Y489 is recognized by multiple hydrophobic interactions in a hydrophobic pocket consisting of VHH residues A50, A57, and A59 (located at or proximal to CDR-2) and I32 (located on CDR-1) and also forms a hydrogen bond with S52 (located on CDR-2) or with the I32 protein backbone (**Figure 3d, 5d**). In this binding mode, this VHH only interacts with a single RBD in trimeric spike. In the two other modes, however, it binds this site but also makes additional interactions with neighboring RBD on another spike protomer.

**Figure 3.**
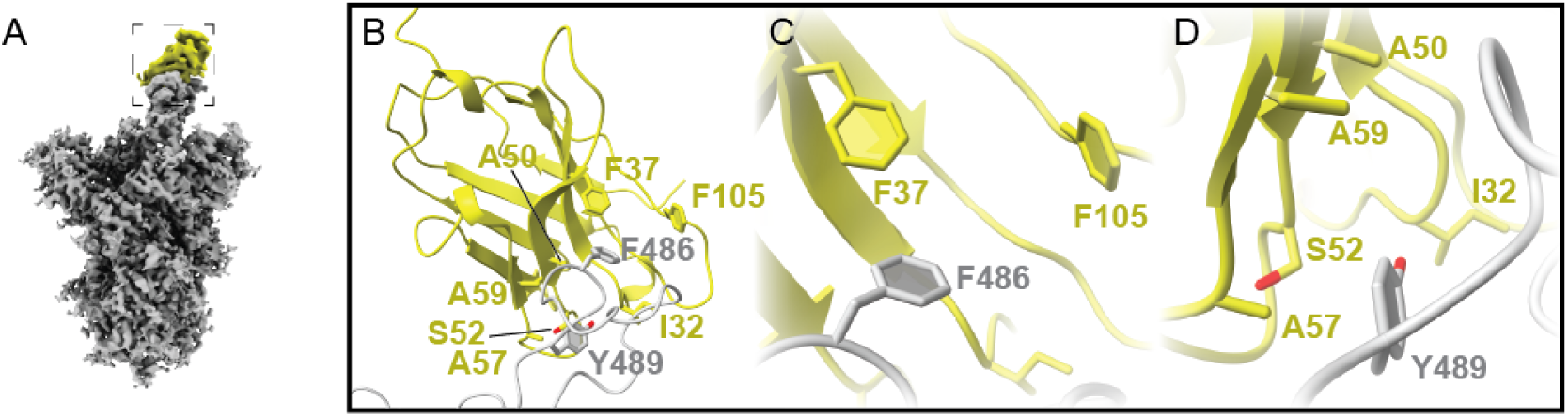
1-1-1 primarily recognizes the peak region of spike RBD. *A*, Overall density map of the 1-1-1 VHH (yellow) bound to SARS-CoV-2 spike protein (grey) RBD in the “up” conformation. Images B, C, and D are located within the dashed box. ***B***, VHH - RBD interacting residues labeled. ***C***, Zoom-in of RBD F486 interactions with VHH residues F37 and F105. F486 most likely forms an aromatic pi-stacking interaction with F37 and F105. ***D***, Zoom-in of RBD Y489 interactions with VHH residues I32, A50, A57, and A59 which form a hydrophobic pocket to stabilize the aromatic Y489. S52 may form a hydrogen bond with Y489 to provide additional stability.

In the second binding mode, the VHH antibody binds the RBD in the “down” conformation (**Figure 4a**). As in the first mode, the VHH recognizes the 485-489 loop (**Figure 4b-d**) but also makes secondary contacts engaging the residues within the RBD-7 epitope class of a neighboring RBD from another protomer if spike trimer (**Figure 4f, Supplementary Figure 1b**). In mode 2, VHH residues G26, G27, and T28 (located on CDR-1) seem to interact with the glycosylation site at N343 of the spike protein (**Figure 4g**). T28 makes an interaction with the acetyl group of the N-acetylglucosamine subunit while G27 and G28 loop around the glycosylation site/group to facilitate binding. VHH F29 (located on CDR-1) rests in an aromatic pocket in the spike protein comprising residues F342, F374, and W436 (**Figure 4h**) and forms pi-stacking interactions with them. A number of individual residue-to-residue interactions also make up this interaction: VHH S30 (located on CDR-1) most likely forms a hydrogen bond with spike S371 (**Figure 4h**), VHH S75 (not located on a CDR) forms a hydrogen bond with spike N440 (**Figure 4i**), and VHH V101 (located on CDR-3) interacts with spike V367 (**Figure 4j**) via hydrophobic interactions. The strength of the map density in this region as well as the strong interactions suggested by the binding data argues that this secondary interaction is playing some role in the recognition of spike by this VHH antibody.

**Figure 4.**
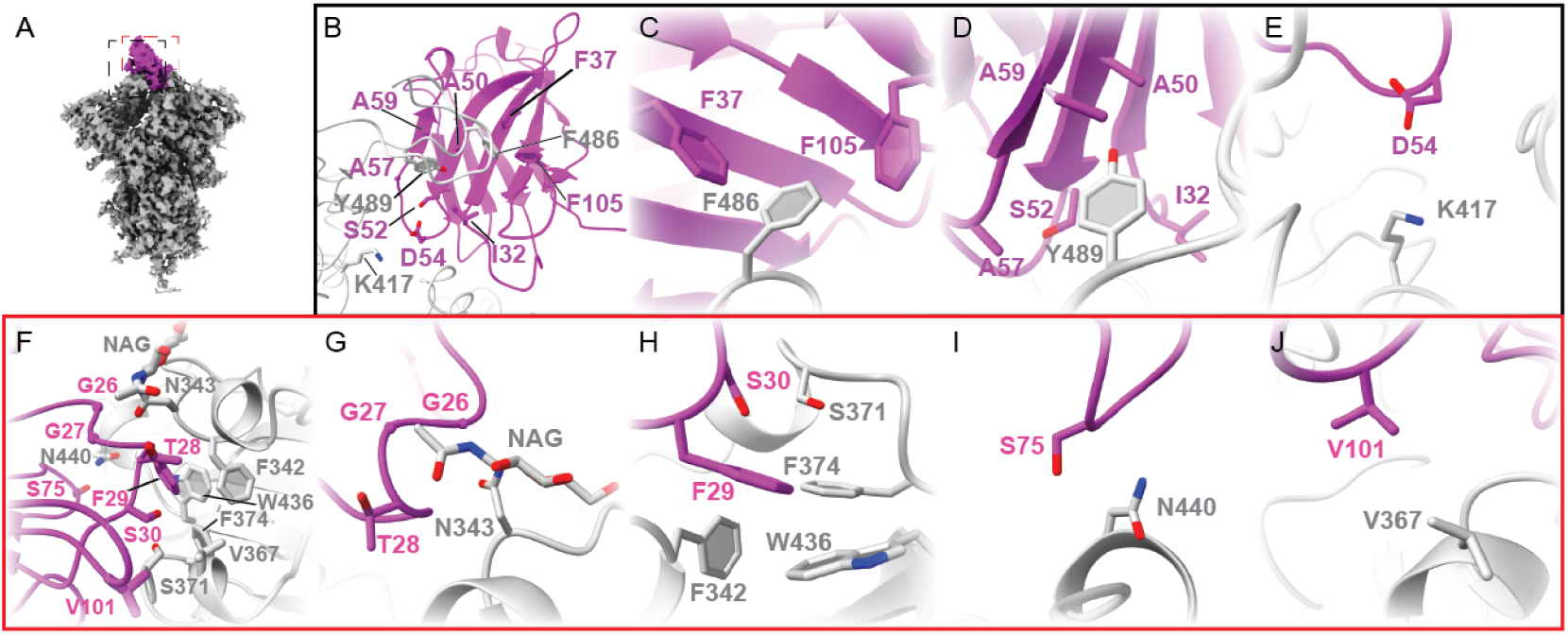
1-1-1 interacts with and recognizes two regions within the spike protein trimer. ***A***, Overall density map of the 1-1-1 VHH (magenta) bound to SARS-CoV-2 spike (gray) RBD in the “down” conformation. Images B, C, D, and E are located within the black dashed box and images F, G, H, I, and J are located on the “backside” of the VHH, within the red dashed box. ***B***, “Frontside” image of VHH and RBD with interacting residues labeled. ***C***, Zoom-in of RBD F486 interactions with VHH residues F37 and F105. Again, F486 most likely forms an aromatic pi-stacking interaction with F37 and F105. ***D***, VHH residues I32, A50, A57, and A59 form a hydrophobic pocket to interact with Y489 and S52, which may form a hydrogen bond with Y489. ***E***, Positively-charged RBD K417 interacts with the negatively-charged VHH residue D54. ***F,*** “Backside” of the VHH-RDB interaction surface (in the “escarpment” region) with interacting residues labeled. ***G***, Loop consisting of G26, G27, and T28 wrapping around glycosylation at RBD 343. G26 and G27 permit the loop to flexibly wrap the glycosylation while T28 may interact with a NAG carbonyl group. ***H***, VHH F29 rests in a hydrophobic, aromatic pocket created by RBD resides F342, F374, and W436 and is most likely stabilized in by multiple pi-stacking interactions. VHH S30 interacts with RBD S371, most likely forming hydrogen bonds with each other. ***I***, VHH S75 interacts with RBD N440, again most likely via hydrogen bonding. ***J,*** VHH V101 makes hydrophobic interactions with RBD V367.

As in mode 1 and mode 2, the VHH in mode 3 recognizes the 485-489 loop (the “peak”) (**Figure 5e-g**) but with different secondary interactions than those seen in mode 2 (**Figure 5i-l, S4c**). The interactions seen here are three individual residue-residue interactions rather than networks of interactions as seen in mode 2, but there is still interaction with residues within the RBD-7 epitope class (RBD-7). Here, VHH E1 (not located on a CDR) interacts with N437 (**Figure 5j**), VHH S25 (located on CDR-1) likely forms a hydrogen bond with Y508 or D405 (**Figure 5k**), and residue T28 (located on CDR-1) interacts with either S275 or S373 (**Figure 5l**). These interactions do not seem to be as strong or defined in the density as the interactions seen in mode 2, suggesting that this surface prefers the “down” RBD conformation in the neighboring RBD.

**Figure 5.**
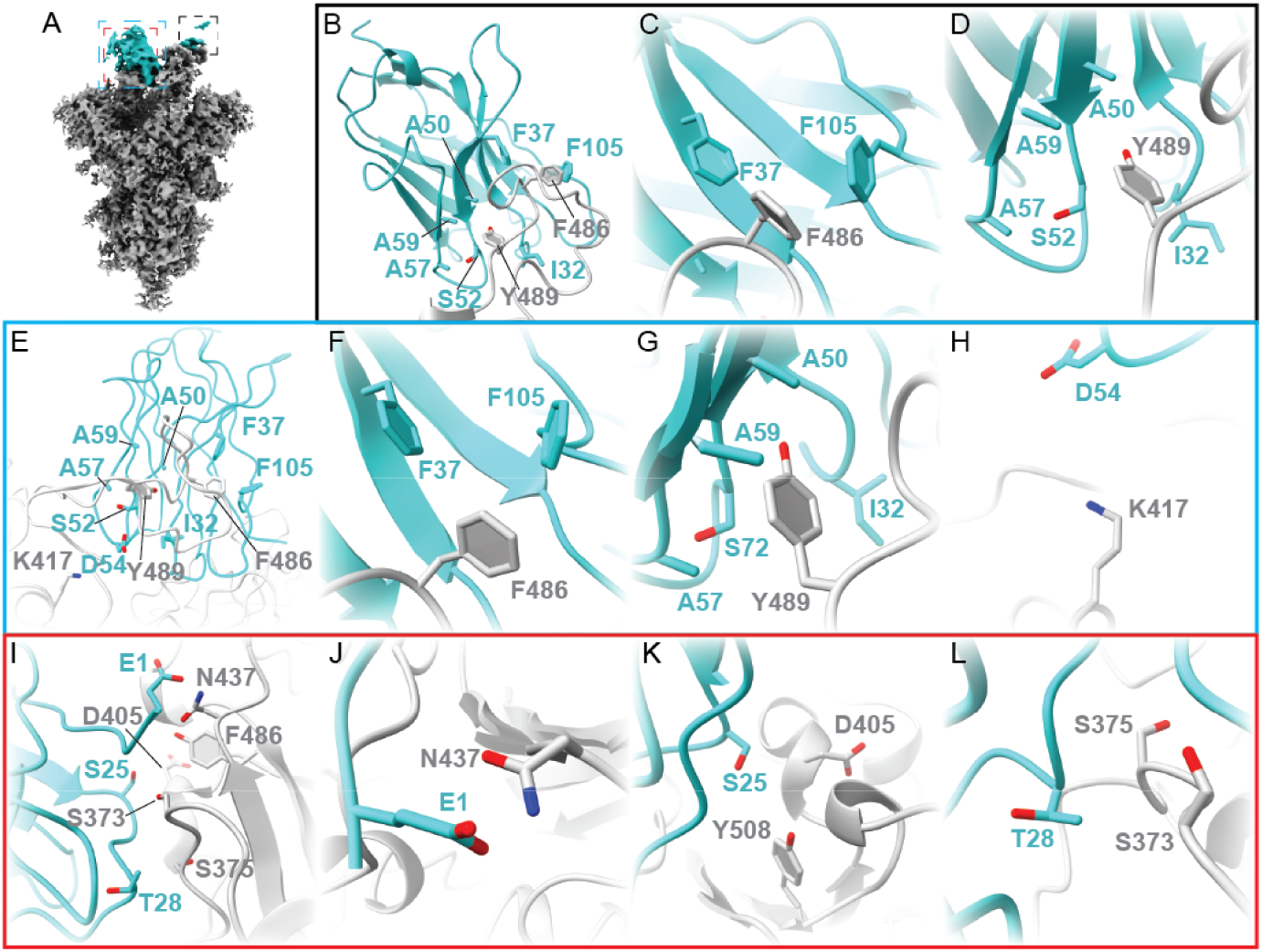
Multiple VHH antibodies can bind the same spike trimer and primarily recognize the “peak” of the RBD. ***A***, Overall density map of two 1-1-1 VHH antibodies (cyan) bound to SARS-CoV-2 spike protein (grey) RBD in the “up” and “down” conformation. Images B, C, and D are located within the black dashed box featuring the VHH on the “up” RBD, images E,F, and G are located on the “frontside” of the VHH on the “down” RBD within the blue dashed box, and images H, I, J, and K are located on the “backside” of the VHH on the “down” RBD within the red dashed box. ***B***, VHH and “up” RBD with interacting residues labeled. ***C***, Zoom-in of RBD F486 interactions with F37 and F105 of the VHH, most likely pi-stacking aromatic interactions ***D***, Zoom-in of RBD Y489 interactions with VHH residues I32, A50, S52, A57, and A59 which form a hydrophobic pocket. ***E***, other VHH and “down” RBD with interacting residues labeled. ***F***, Zoom-in of RBD F486 interactions with F37 and F105 of the VHH. ***G***, Zoom-in of RBD Y489 interactions with VHH residues I32, A50, A57, and A59 which also form a hydrophobic pocket and S52 which may form a hydrogen bond with Y489 ***H***, zoom in of RBD K417 interacting with VHH D54. Most likely, these residues interact via their opposite charges. ***I,*** “Backside” interaction surface of VHH with RBD with interacting residues labeled. This surface is different from the surface seen in class 2 and seems to be a much weaker interaction, which can be explained by the associated RBD being in the “up” rather than the “down” conformation. This suggests that this recognition of the spike protein by the VHH “backside” plays a secondary role in comparison to the interactions seen in the “frontside.” ***J***, VHH E1 interacts with RBD N437, ***K***, VHH S25 interacts with either RBD Y508 and D405, and ***L***, VHH T28 interacts with either RBD S373 and S375-all of which are most likely hydrogen bonding interactions.

Previously, K417 was identified as a key residue in TB201-1’s binding epitope^24^. In our structures, residue D54 of the VHH was found in close proximity to K417 of the RBD in mode 2 and 3 (**Figure 4e, 5h**). When the VHH is bound to the RBD in the “up” conformation (mode 1), these two residues are separated by a much larger distance which may hinder their interaction. Given this close distance and the fact that mutation of K417 affects the ability of TB201-1 to recognize spike protein^24^, K417 recognition may be important for VHH recognition of the RBD while in the “down” conformation but does not play a role in recognition while in the “up” conformation. The other residues identified previously (F456, G476, and N487 of the RBD) have no obvious interaction surface/residues with the VHH and may play a lesser role in recognition^24^. The VHH antibodies or components observed in the maps may be part of the same VHH-Fc trimer or from separate molecules since an unresolved eleven-residue linker connects the Fab regions of the Fc domain.

The commonality of the binding site at the RBD-1 epitope in our structural data and the binding data which shows that TB201-1 (monomer) has an increase in binding for S1 trimer over S1 monomer suggests that the RBD-1 epitope is the primary binding site of this nanobody while the other set of interactions (at RBD-7) is a secondary binding site that assists the nanobody in binding the RBD when in the “down” position (**Figure 1b**).

## Discussion

We engineered synthetic multivalent VHH-Fc antibodies by leveraging a series of high-throughput technologies, including large-scale DNA synthesis and assembly (to fabricate antibody libraries), phage display (to screen the libraries), long-read NGS (to evaluate trimeric permutations of VHH sequences), and biophysical assays (to triage leads). The combination of these approaches yielded hexavalent anti-S1 VHH-Fc antibodies that outperformed their parental bivalent VHH-Fc constructs in binding and competition assays incorporating a variety of S variants. The antibodies described here add to the growing armamentarium of SARS-CoV-2 antibodies that have been discovered since the onset of the COVID-19 pandemic. Additionally, we have characterized the binding of VHH antibody 1-1-1 to SARS-CoV-2 spike protein trimer. Here, we identify the residues critical for spike recognition by the VHH and use this knowledge to improve our understanding of how various SARS-CoV-2 variants evade binding by this VHH.

Multivalent antibodies are a more recent addition to this armamentarium against SARS-CoV-2. A few groups have capitalized on antibody structure and epitope information to engineer human-derived IgG1-like bispecific antibodies,^15^ multivalent VHH fusions,^8^ dual variable domain Igs,^14^ tetravalent VHH-p53 fusions,^27^ and multivalent VHH-apoferritin fusions^28^ from relatively small collections of characterized mAbs. The remainder have largely used phage display to initially screen for leads, from which they engineered multivalent constructs.^11,12,13,16,17,30,31^ In aggregate, these efforts yielded numerous potently neutralizing anti-S1 antibodies, many of which retained this activity against SARS-CoV-2 variants of concern, including the Alpha, Beta, Gamma, and Delta variants. These studies also identified multiple unique binding and neutralization mechanisms. To provide a few examples, Cho and colleagues^14^ generated a bispecific antibody capable of simultaneously binding the receptor-binding domain (RBD) and N-terminal domain (NTD) epitopes of S1; Lim and colleagues^16^ demonstrated that combining neutralizing and non-neutralizing antibody fragments into bispecific VH/Fab IgGs could improve neutralizing activity; Koenig and colleagues^8^ found that targeting bispecifics to two independent vulnerable RBD epitopes could minimize viral escape; and Wu and colleagues^31^ showed that two copies of a RBD-specific VHH could be multimerized with a VHH specific to human serum albumin to improve *in vivo* efficacy by both increasing neutralizing activity and half-life. Although monospecific constructs are poorly represented among the multivalent anti-SARS-CoV-2 antibodies discovered so far, the few reports describing them indicate that increasing the avidity of a monospecific antibody through multimerization is sufficient to boost its therapeutic efficacy.^28,29,31^

Where our approach departs from others is in the use of an unbiased method for screening multivalents. By combining phage display with long-read amplicon sequencing, we screened thousands of trimeric permutations to empirically determine where distinct VHH fragment sequences performed best within the trimeric assembly. Although TB201-1 was the most frequent VHH fragment in all three positions of the tandem assembly, it was more frequently found in the secondary than the primary and tertiary positions. The second most frequently observed VHH fragment, TB202-76, showed the opposite trend, instead preferring the primary and tertiary positions. As mentioned above, TB202-76 cross-competes with TB201-1 for S1 binding and, like TB201-1, blocks the S1-ACE2 interaction and neutralizes authentic SARS-CoV-2.^24^ Nevertheless, it is surprising that TB202-76 appeared so frequently in the enriched trimeric phage libraries given its middling biophysical characteristics relative to the other VHH fragments identified from our synthetic libraries.^24^ Many of the monovalent antibodies used in this work — TB202-63, TB201-20, TB201-34, TB201-1, and TB202-3 — target overlapping epitopes on the outer face of RBD (described as the RBD-4 community in Hastie et al.^23^).^24^ As such, these antibodies likely competed with one another during biopanning, with binding orientation, the extent of epitope overlap, inter-VHH distance, and ability to cross-link S1 proteins potentially playing roles in determining which tandem VHH arrangements are enriched during biopanning. The former two factors (binding orientation and epitope overlap) may have also contributed to the better performance of 76-1-1 versus 1-1-1 in our S1-ACE2 competition assay but not SPR. Additionally, the latter two factors (inter-VHH distance and ability to cross-link S1 proteins) may explain why TB202-76 was able to bind the Omicron S trimer when 76-1-1 was not, as the TB202-76 domains in 76-1-1 are spaced further apart than those in TB202-76. Further work is needed, however, to confirm these possibilities.

In our structural studies, we identified a primary VHH epitope on the spike protein located on the RBD-1 region that was common to all structural classes determined. The RBD-1 region is involved in ACE2 interactions and is often mutated in SARS-CoV-2 variants. Some of these mutations are favored as they increase cell-cell fusion via increased ACE2 binding affinity^34^ while others contribute to immune evasion of potent antibodies^35^ and several of these variant mutations could adversely affect binding with the VHH antibodies studied here. RBD residue F486, which is one of the primary residues recognized by the VHH antibody, was mutated to valine in a few Omicron strains (**Figure 3c**). Although this mutation allows the residue to maintain its hydrophobic character, the loss of its aromatic nature oblates the pi-stacking interaction with VHH residues F37 and F105 explaining the observed loss of binding. VHH binding is also sensitive to the K417N mutation as it abrogates the interaction between positively-charged spike residue K417 and negatively-charged VHH residue D54 (**Figure 4e**). The Q498R spike mutation may generate a steric clash to prevent close interaction with the VHH. Collectively, these mutations may result in the observed loss of VHH binding.

The VHH’s secondary interaction surface aligns with the RBD-7 epitope and the residues involved in interaction on that epitope differ depending on the conformation of the spike trimer. In the class 2 structure, mutation G339D may deform the loop where the glycosylated N343 residue is located and may make VHH recognition of that site more difficult (**Figure 4g**). Additionally, mutations S371F, S373P and S375F all alter the loop conformation at this region and thus interfere with VHH recognition. Mutation N440K may become too bulky to accommodate its interaction with VHH S75 **(Figure 4i**). In the class 3 secondary interaction surface, mutation S373P and S375F would prevent hydrogen bonds of VHH T28 with spike S373 and S375 (**Figure 5l**).

In conclusion, we improved the affinity of anti-S antibodies against SARS-CoV-2 variants by trimerizing the VHH sequences of VHH-Fc antibodies. We also developed a high-throughput approach for screening multivalent permutations of VHH fragments. Unlike existing approaches for engineering multivalents, our approach identifies the positional preferences of each screened VHH fragment in an unbiased manner. As a result, this approach may yield further insights into the mechanisms through which multivalency improves the avidity and, in turn, the diagnostic or therapeutic performance of VHH antibodies.

## Materials and Methods

### Cloning

The VHH-Fc antibodies (TB201-1, TB202-76, TB202-85, and so on) described in this paper were created by reformatting a series of VHH single-domain antibodies discovered by phage display.^24^ This process involved synthesizing back-translated VHH protein sequences and cloning them into the mammalian fc-fusion IgG1 expression vector pTwist_CMV_BG_WPRE_Neo-VHH-Fc. The resulting DNA constructs were obtained as purified, transfection-ready plasmid DNA using the Twist eCommerce Portal. VHH-Fc antibodies were produced in HEK Expi293 cells (Thermo Fisher Scientific) and purified using Protein A resin (PhyNexus) and the Hamilton Microlab STAR platform. Antibody quality control was performed using capillary electrophoresis sodium dodecyl sulfate (CE-SDS).

Hexavalent VHH-Fc antibodies were generated by assembling trimeric VHH sequences and then inserting these assemblies into a backbone vector. Internal assembly sites (those connecting VHH sequences with one another) were designed to include Type II BsaI restriction sites (for directional cloning of trimeric VHH assemblies), whereas external assembly sites (those connecting the trimer VHH assembly to the backbone vector) were designed to include homologous overhaFngs for insertion into a fc-fusion IgG1 acceptor vector (Figure 1a) or the pADL-22c phage vector (Figure 2a). Internal assembly sites were incorporated into the (GGGGS)_2_AS linker sequences that were inserted in between each VHH sequence. VHH sequences with cloning flanks were ordered as gene fragments using the Twist eCommerce Portal.

### Phage display

The phage library (Figure 2a) was electroporated into TG1 *E. coli* cells (Lucigen) for propagation. Phage particles were blocked with phosphate buffered saline (PBS) with 5% bovine serum albumin (BSA) and depleted for non-specific binders on M-280 streptavidin coated magnetic beads (Thermo). Biotinylated SARS-CoV-2 S1 protein was mixed with M-280 beads, washed with PBS/0.5% Tween-20 to remove unbound protein and used as panning target for four rounds of panning for final antigen concentrations of 100 pM for Round 1 and 50 pM for Round 2. Phage supernatants depleted of non-specific binders were transferred to a bead mixture containing bound biotinylated SARS-CoV-2 S1 and allowed to bind for 1 hour at RT to select for binders with gentle rotation. Following incubation, beads were washed several times with PBS/0.5% Tween-20 to remove non-binding clones. Remaining bound phage were eluted with trypsin in PBS buffer for 30 minutes at 37°C. The output supernatant enriched in binding clones was amplified in TG1 *E. coli* cells to use as input phage for the next round of selection, with each round increasing the wash cycles and lowering the total amount of antigen present.

Bacterial colonies containing the phagemid display vector were lawned on 2YT agar plates containing 100 µg/mL carbenicillin. After an overnight incubation, 2YT media with 100 µg/mL of carbenicillin was added to the lawned plate and output was scrapped and collected; glycerol was added to a final concentration of 25% to generate glycerol stocks. 100 µL of round output glycerol stock was miniprepped in preparation for NGS sequencing. Round output was amplified (25 cycles) using primers “LoopGenomics_Amp”, “LoopGenomics_Amp_R” (ATGTTGTGTGGAATTGTGAGCGGAT, ACAGCCCTCATAGTTAGCGTAACGA), which bind to padl22-c backbone, and the Q5 Polymerase 2X Master Mix. Amplified round output was size-verified using the BioAnalyzer High Sensitivity DNA Analysis Kit. Sequencing libraries were prepared from long range PCR products with the commercially available LoopSeq kits from Loop Genomics (protocols available at https://www.loopgenomics.com/product-page/loopseq-pcr-amplicon-3x8-plex-kit). The process involves attaching a barcoding adapter containing both a Unique Molecular Identifier (UMI) and a sample index to each molecule in the same sample. Barcoded molecules were amplified, multiplexed, and each UMI distributed intramolecularly to a random position within each parent molecule. Molecules were then fragmented into smaller units, creating a library of UMI-tagged fragments with an average length of 400 bp. After sequencing this library on a NextSeq machine, synthetic long reads were constructed from the short-read sequencing reads using the standard Loop Genomics informatics pipeline. VHH monomer frequency was analyzed per trimer position.

### SPR

Surface plasmon resonance (SPR) experiments were performed on a Carterra LSA SPR biosensor equipped with a HC30M chip at 25°C in HBS-TE (10 mM HEPES pH 7.4, 150 mM NaCl, 3 mM EDTA, 0.05% Tween-20). SARS-CoV-2 S glycoproteins were diluted to 5 μg/mL and amine-coupled to the sensor chip by EDC/NHS activation, followed by ethanolamine HCl quenching. Increasing concentrations of antibody were flowed over the sensor chip in HBS-TE with 0.5 mg/mL BSA with 5 minute association and 15 minute dissociation. Following each injection cycle, the surface was regenerated with 2x 30 second injections of IgG elution buffer (Thermo Fisher Scientific). Data were analyzed in Carterra’s Kinetics Tool software with 1:1 binding model.

### Sandwich ELISA

For the sandwich ELISA, TB201-1 Fc or 1-1-1 were immobilized in 96-well plates and blocked with PBS supplemented with 0.5% (w/v) BSA. Twelve-point serial dilutions (1:3) of prefusion-stabilized SARS-CoV-2 S trimers were prepared for each of the tested S variants, all with 24 nM starting concentrations. Biotinylated TB202-24 (0.4 μg/mL), an antibody that does not compete with TB201-1 for S binding, and streptavidin-HRP (1:5,000) were used for detection. All ELISAs were performed in quadruplicate (n=4).

### Flow Cytometry Competition Assay

Antibodies were also assayed by flow cytometry to measure inhibition of S1 binding to Vero E6 cells, which constitutively express ACE2. S1-RBD-mFc was purchased from Acro Biosystems; other S1 variant constructs were designed and ordered as transfection-ready clonal genes in the pTwist_CMV_BG_WPRE_Neo base mammalian expression vector using the Twist eCommerce Portal., transfected in HEK293 cells, and the resulting proteins purified using Protein A resin (PhyNexus) and the Hamilton Microlab STAR platform. For the inhibition assay, Vero E6 cells were plated in 96-well plates at 1.5 x 10^5^ cells per well. Antibodies were serially diluted 1:3 in PBS from a 100 nM starting concentration. These dilutions were then incubated with 0.1 µg/mL S1-RBD-mFc (or analogous variant protein) at 4° C for 1 hour. The antibody-S1-RBD-mFc (or analogous variant protein) mixture was then added to the plated Vero E6 cells and incubated at 4° C for 1 hour. After three washes with PBS, the plated cells were incubated with an allophycocyanin (APC)-conjugated anti-mouse antibody for 1 hour at 4° C. Cells were analyzed by flow cytometry by measuring the APC signal.

### Spike protein expression and purification

SARS-CoV-2 Spike-6P trimer protein carrying WA.1 was expressed and purified by transfecting Expi293F cells using WA.1-spike-6P plasmids as described previously^36^. Transfections were performed as per the manufacturer’s protocol (Thermo Fisher). Briefly, Expi293F cells (2.5 x 10^6^ cells/mL) were transfected using ExpiFectamine^TM^ 293 transfection reagent (ThermoFisher, cat. no. A14524). The cells were harvested 4-5 days post-transfection. The spike protein was purified using His-Pur Ni-NTA affinity purification method. The column was washed with Buffer containing 25 mM Imidazole, 6.7 mM NaH_2_PO_4_·H_2_O and 300 mM NaCl in PBS followed by spike protein elution in elution buffer containing 235 mM Imidazole, 6.7 mM NaH_2_PO_4_•H_2_O and 300 mM NaCl in PBS. Eluted protein was dialyzed against PBS and concentrated. The concentrated protein was loaded onto a Superose-6 Increase 10/300 column and protein eluted as trimeric spike collected. Protein quality was evaluated by SDS-PAGE and by Negative Stain-EM.

### Negative Stain – Electron Microscopy (NS-EM)

Spike protein was diluted to 0.05 mg/mL in PBS before grid preparation. A 3 μL drop of diluted protein (∼0.025 mg/mL) was applied to previously glow-discharged, carbon-coated grids for ∼60 seconds, blotted and washed twice with water, stained with 0.75% uranyl formate, blotted and air-dried. Between 30 and 50 images were collected on a Talos L120C microscope (Thermo Fisher) at 73,000 magnification and 1.97 Å pixel size. Relion-3.1^37^ or Cryosparc v3.3.2^38^ was used for particle picking and 2D classification.

### Sample preparation for Cryo-EM

SARS-CoV-2 spike-6P trimer at 0.6 mg/mL concentration was incubated with a 0.4 sub-molar ratioVHH to prevent inter-spike cross linking, mediated by bivalent binding of intact antibody. The complex was incubated at RT for ∼5 minutes before vitrification. Three μL of the complex was applied onto a freshly glow-discharged (PLECO easiGLOW) 400 mesh, 1.2/1.3 C-Flat grid (Electron Microscopy Sciences). After 20 seconds of incubation, grids were blotted for 3 seconds at 0 blot force and vitrified using a Vitrobot IV (Thermo Fisher Scientific) under 22°C with 100% humidity.

### Cryo-EM data acquisition

Single-particle Cryo-EM data for WA.1 spike-nanobody complexes of VHH TB201-1 were collected on a 300 kV Titan Krios transmission electron microscope (ThermoFisher Scientific) equipped with Gatan K3 direct electron detector behind 30 eV slit width energy filter (at the National Center for CryoEM Access and Training). Multi-frame movies were collected at a pixel size of 1.0691 Å per pixel with a total dose of 63.81 e/Å_2_ at defocus range of -0.6 to -2.6 μm.

### Cryo-EM data analysis and model building

Cryo-EM movies were motion-corrected by Patch motion correction implemented in Cryosparc v3.3.1^38^. Motion-corrected micrographs were corrected for contrast transfer function using Cryosparc’s implementation of Patch CTF estimation. Micrographs with poor CTF fits were discarded using CTF fit resolution cutoff to ∼6.0 Å. Particles were picked using a Blob picker, extracted and subjected to an iterative round of 2D classification (**Supplementary Figure 2A**). Particles belonging to the best 2D classes with secondary structure features were selected for heterogeneous 3D refinement to separate VHH-bound spike particles from non-VHH bound spike particles **(Supplementary Figure 2B**). Particles belonging to the best VHH-bound 3D classes were refined in non-uniform 3D refinement with per particle CTF and higher-order aberration correction (**Supplementary Figure 2C,D**).

To further improve the resolution of the RBD-VHH binding interface, a soft mask was created covering the VHH and the proximal spike surface and refined the map locally in Cryosparc using Local Refinement on signal subtracted particles (**Supplementary Figure 2E**). All maps were density modified in Phenix^39^ using Resolve Cryo-EM. The combined Focused Map tool in Phenix was used to integrate high resolution locally refined maps into an overall map. Additional data processing details are summarized in **Supplementary Tables 1, 2, and 3**. The initial spike models for WA.1 (PDB:7lrt) as well as VHH (generated with Alphafold^40^) were docked into the combined focused Cryo-EM density maps using UCSF ChimeraX^41^. The full Spike-VHH model was refined using rigid body refinement in Phenix and refined further using real-space refinement. Glycosylations with visible density were modeled in Coot^42^. Model validation was performed using Molprobity^43^. Figures were prepared in ChimeraX.

## Abbreviations

ACE2: angiotensin-converting enzyme 2
APC: allophycocyanin
BSA: bovine serum albumin
CDR: complementarity-determining region
CE-SDS: capillary electrophoresis sodium dodecyl sulfate
COVID-19: coronavirus disease 2019
Fab: fragment antigen-binding
HRP: horseradish peroxidase
Ig(G): immunoglobulin (G)
NGS: next-generation sequencing
NTD: N-terminal domain
PBS: phosphate-buffered saline
RBD: receptor-binding domain
RT: room temperature
RT-PCR: real-time
PCR S: spike
SARS-CoV-2: severe acute respiratory coronavirus 2
scFv: single-chain variable fragment
SPR: surface plasmon resonance
UMI: unique molecular identifier
VHH: variable domain of camelid heavy-chain

## Acknowledgements

Research reported in this publication was supported by the National Institute of Biomedical Imaging and Bioengineering of the National Institutes of Health (under award numbers 75N92019P00328, U54EB015408, and U54EB027690) as part of the Rapid Acceleration of Diagnostics (RADx) initiative, launched to speed innovation in the development, commercialization, and implementation of technologies for COVID-19 testing. The funders had no role in the decision to submit the work for publication and the views expressed herein are the authors’ and do not necessarily represent the views of the National Institutes of Health or the United States Department of Health and Human Services. E.A.O. was supported by the NIDDK award 5R01DK115213. The Titan Krios Cryo-EM data sets were collected at the National Center for CryoEM Access and Training (NCCAT) and the Simons Electron Microscopy Center located at the New York Structural Biology Center, supported by the NIH Common Fund Transformative High Resolution Cryo-Electron Microscopy program (U24 GM129539,) and by grants from the Simons Foundation (SF349247) and NY State Assembly. We also thank the staff of Robert P. Apkarian Integrated Electron Microscopy Core (IEMC) at Emory University, Atlanta for their support with preliminary sample screening on Talos Arctica.

## References

1. Batista FD, Neuberger MS. Affinity dependence of the B cell response to antigen: a threshold, a ceiling, and the importance of off-rate. Immunity 1998; 8:751–9.

2. Bradbury ARM, Sidhu S, Dübel S, McCafferty J. Beyond natural antibodies: the power of in vitro display technologies. Nature Biotechnology 2011; 29:245–54. Available from: 10.1038/nbt.1791

3. Kang TH, Seong BL. Solubility, Stability, and Avidity of Recombinant Antibody Fragments Expressed in Microorganisms. Front Microbiol 2020; 11:1927.

4. Czajka TF, Vance DJ, Mantis NJ. Slaying SARS-CoV-2 One (Single-domain) Antibody at a Time. Trends Microbiol 2021; 29:195–203.

5. Schoof M, Faust B, Saunders RA, Sangwan S, Rezelj V, Hoppe N, Boone M, Billesbølle CB, Puchades C, Azumaya CM, et al. An ultrapotent synthetic nanobody neutralizes SARS-CoV-2 by stabilizing inactive Spike. Science 2020; 370:1473–9.

6. Wrapp D, De Vlieger D, Corbett KS, Torres GM, Wang N, Van Breedam W, Roose K, van Schie L, VIB-CMB COVID-19 Response Team, Hoffmann M, et al. Structural Basis for Potent Neutralization of Betacoronaviruses by Single-Domain Camelid Antibodies. Cell 2020; 181:1004–15.e15.

7. Huo J, Le Bas A, Ruza RR, Duyvesteyn HME, Mikolajek H, Malinauskas T, Tan TK, Rijal P, Dumoux M, Ward PN, et al. Neutralizing nanobodies bind SARS-CoV-2 spike RBD and block interaction with ACE2. Nat Struct Mol Biol 2020; 27:846–54.

8. Koenig P-A, Das H, Liu H, Kümmerer BM, Gohr FN, Jenster L-M, Schiffelers LDJ, Tesfamariam YM, Uchima M, Wuerth JD, et al. Structure-guided multivalent nanobodies block SARS-CoV-2 infection and suppress mutational escape. Science 2021; 371. Available from: 10.1126/science.abe6230

9. Xiang Y, Nambulli S, Xiao Z, Liu H, Sang Z, Duprex WP, Schneidman-Duhovny D, Zhang C, Shi Y. Versatile and multivalent nanobodies efficiently neutralize SARS-CoV-2. Science 2020; 370:1479–84.

10. Hanke L, Vidakovics Perez L, Sheward DJ, Das H, Schulte T, Moliner-Morro A, Corcoran M, Achour A, Karlsson Hedestam GB, Hällberg BM, et al. An alpaca nanobody neutralizes SARS-CoV-2 by blocking receptor interaction. Nat Commun 2020; 11:4420.

11. Dong J, Huang B, Wang B, Titong A, Gallolu Kankanamalage S, Jia Z, Wright M, Parthasarathy P, Liu Y. Development of humanized tri-specific nanobodies with potent neutralization for SARS-CoV-2. Sci Rep 2020; 10:17806.

12. Dong J, Huang B, Jia Z, Wang B, Gallolu Kankanamalage S, Titong A, Liu Y. Development of multi-specific humanized llama antibodies blocking SARS-CoV-2/ACE2 interaction with high affinity and avidity. Emerg Microbes Infect 2020; 9:1034–6.

13. Lu Q, Zhang Z, Li H, Zhong K, Zhao Q, Wang Z, Wu Z, Yang D, Sun S, Yang N, et al. Development of multivalent nanobodies blocking SARS-CoV-2 infection by targeting RBD of spike protein. J Nanobiotechnology 2021; 19:33.

14. Cho H, Gonzales-Wartz KK, Huang D, Yuan M, Peterson M, Liang J, Beutler N, Torres JL, Cong Y, Postnikova E, et al. Bispecific antibodies targeting distinct regions of the spike protein potently neutralize SARS-CoV-2 variants of concern. Sci Transl Med 2021; 13:eabj5413.

15. De Gasparo R, Pedotti M, Simonelli L, Nickl P, Muecksch F, Cassaniti I, Percivalle E, Lorenzi JCC, Mazzola F, Magrì D, et al. Bispecific IgG neutralizes SARS-CoV-2 variants and prevents escape in mice. Nature 2021; 593:424–8.

16. Lim SA, Gramespacher JA, Pance K, Rettko NJ, Solomon P, Jin J, Lui I, Elledge SK, Liu J, Bracken CJ, et al. Bispecific VH/Fab antibodies targeting neutralizing and non-neutralizing Spike epitopes demonstrate enhanced potency against SARS-CoV-2. MAbs 2021; 13:1893426.

17. Bracken CJ, Lim SA, Solomon P, Rettko NJ, Nguyen DP, Zha BS, Schaefer K, Byrnes JR, Zhou J, Lui I, et al. Bi-paratopic and multivalent VH domains block ACE2 binding and neutralize SARS-CoV-2. Nat Chem Biol 2021; 17:113–21.

18. Moliner-Morro A, J Sheward D, Karl V, Perez Vidakovics L, Murrell B, McInerney GM, Hanke L. Picomolar SARS-CoV-2 Neutralization Using Multi-Arm PEG Nanobody Constructs. Biomolecules 2020; 10. Available from: 10.3390/biom10121661

19. Hunt AC, Case JB, Park Y-J, Cao L, Wu K, Walls AC, Liu Z, Bowen JE, Yeh H-W, Saini S, et al. Multivalent designed proteins protect against SARS-CoV-2 variants of concern. bioRxiv 2021; Available from: 10.1101/2021.07.07.451375

20. Muyldermans S. A guide to: generation and design of nanobodies. FEBS J 2021; 288:2084–102.

21. Winter G, Milstein C. Man-made antibodies. Nature 1991; 349:293–9.

22. Benhar I. Design of synthetic antibody libraries. Expert Opin Biol Ther 2007; 7:763–79.

23. Hastie KM, Li H, Bedinger D, Schendel SL, Dennison SM, Li K, Rayaprolu V, Yu X, Mann C, Zandonatti M, et al. Defining variant-resistant epitopes targeted by SARS-CoV-2 antibodies: A global consortium study. Science 2021; 374:472–8.

24. Yuan TZ, Garg P, Wang L, Willis JR, Kwan E, Hernandez AGL, Tuscano E, Sever EN, Keane E, Soto C, et al. Rapid discovery of diverse neutralizing SARS-CoV-2 antibodies from large-scale synthetic phage libraries. MAbs 2022; 14:2002236.

25. Kosuri S, Eroshenko N, Leproust EM, Super M, Way J, Li JB, Church GM. Scalable gene synthesis by selective amplification of DNA pools from high-fidelity microchips. Nat Biotechnol 2010; 28:1295–9.

26. Yuan TZ, Garg P, Willis JR, Kwan E, Lujan Hernandez AG, Tuscano E, Sever EN, Keane E, Soto C, Mucker EM, Fouch ME, Davidson E, Doranz BJ, Kailasan S, Aman MJ, Li H, Hooper JW, Saphire EO, Crowe Jr JE, Liu Q, Axelrod F, and Sato AK. Rapid discovery of diverse neutralizing SARS-CoV-2 antibodies from large-scale synthetic phage libraries. MAbs in review.

27. Leach A, Miller A, Bentley E, Mattiuzzo G, Thomas J, McAndrew C, Van Montfort R, Rabbitts T. Implementing a method for engineering multivalency to substantially enhance binding of clinical trial anti-SARS-CoV-2 antibodies to wildtype spike and variants of concern proteins. Sci Rep 2021; 11:10475.

28. Rujas E, Kucharska I, Tan YZ, Benlekbir S, Cui H, Zhao T, Wasney GA, Budylowski P, Guvenc F, Newton JC, et al. Multivalency transforms SARS-CoV-2 antibodies into ultrapotent neutralizers. Nat Commun 2021; 12:3661.

29. Xu J, Xu K, Jung S, Conte A, Lieberman J, Muecksch F, Lorenzi JCC, Park S, Schmidt F, Wang Z, et al. Nanobodies from camelid mice and llamas neutralize SARS-CoV-2 variants. Nature 2021; 595:278–82.

30. Miersch S, Li Z, Saberianfar R, Ustav M, Brett Case J, Blazer L, Chen C, Ye W, Pavlenco A, Gorelik M, et al. Tetravalent SARS-CoV-2 Neutralizing Antibodies Show Enhanced Potency and Resistance to Escape Mutations. J Mol Biol 2021; 433:167177.

31. Wu X, Cheng L, Fu M, Huang B, Zhu L, Xu S, Shi H, Zhang D, Yuan H, Nawaz W, et al. A potent bispecific nanobody protects hACE2 mice against SARS-CoV-2 infection via intranasal administration. Cell Rep 2021; 37:109869.

32. Jovčevska, I., & Muyldermans, S. (2020). The Therapeutic Potential of Nanobodies. BioDrugs : clinical immunotherapeutics, biopharmaceuticals and gene therapy, 34(1), 11–26. 10.1007/s40259-019-00392-z

33. Hastie KM, Li H, Bedinger D, Schendel SL, Dennison SM, Li K, Rayaprolu V, Yu X, Mann C, Zandonatti M, et al. Defining variant-resistant epitopes targeted by SARS-CoV-2 antibodies: A global consortium study. Science 2021; 374:472–8.

34. Tian D, Sun Y, Xu H, Ye Q. The emergence and epidemic characteristics of the highly mutated SARS-CoV-2 Omicron variant. J Med Virol. 2022 Jun;94(6):2376–2383. doi: 10.1002/jmv.27643. Epub 2022 Feb 11. PMID: 35118687; PMCID: PMC9015498.

35. Yuan TZ, Garg P, Wang L, Willis JR, Kwan E, Hernandez AGL, Tuscano E, Sever EN, Keane E, Soto C, et al. Rapid discovery of diverse neutralizing SARS-CoV-2 antibodies from large-scale synthetic phage libraries. MAbs 2022; 14:2002236.

36. Kumar S, Patel A, Lai L, Chakravarthy C, Valanparambil R, Reddy ES, Gottimukkala K, Davis-Gardner ME, Edara VV, Linderman S, Nayak K, Dixit K, Sharma P, Bajpai P, Singh V, Frank F, Cheedarla N, Verkerke HP, Neish AS, Roback JD, Mantus G, Goel PK, Rahi M, Davis CW, Wrammert J, Godbole S, Henry AR, Douek DC, Suthar MS, Ahmed R, Ortlund E, Sharma A, Murali-Krishna K, Chandele A. Structural insights for neutralization of Omicron variants BA.1, BA.2, BA.4, and BA.5 by a broadly neutralizing SARS-CoV-2 antibody. Sci Adv. 2022 Oct 7;8(40):eadd2032. doi: 10.1126/sciadv.add2032. Epub 2022 Oct 5. PMID: 36197988; PMCID: PMC9534492.

37. S. H. Scheres, RELION: implementation of a Bayesian approach to cryo-EM structure determination. J Struct Biol 180, 519–530 (2012).

38. A. Punjani, J. L. Rubinstein, D. J. Fleet, M. A. Brubaker, cryoSPARC: algorithms for rapid unsupervised cryo-EM structure determination. Nat Methods 14, 290–296 (2017).

39. P. D. Adams et al., PHENIX: a comprehensive Python-based system for macromolecular structure solution. Acta Crystallogr D Biol Crystallogr 66, 213–221 (2010).

40. Jumper, J. et al. “Highly accurate protein structure prediction with AlphaFold.” Nature, 596, pages 583–589 (2021). DOI: 10.1038/s41586-021-03819-2

41. T. D. Goddard et al., UCSF ChimeraX: Meeting modern challenges in visualization and analysis. Protein Sci 27, 14–25 (2018).

42. P. Emsley, B. Lohkamp, W. G. Scott, K. Cowtan, Features and development of Coot. Acta Crystallogr D Biol Crystallogr 66, 486–501 (2010).

43. C. J. Williams et al., MolProbity: More and better reference data for improved all-atom structure validation. Protein Sci 27, 293–315 (2018).

