## Supplementary Materials for "Structural basis of SARS-Cov-2 spike recognition by engineered synthetic multivalent VHH antibodies"

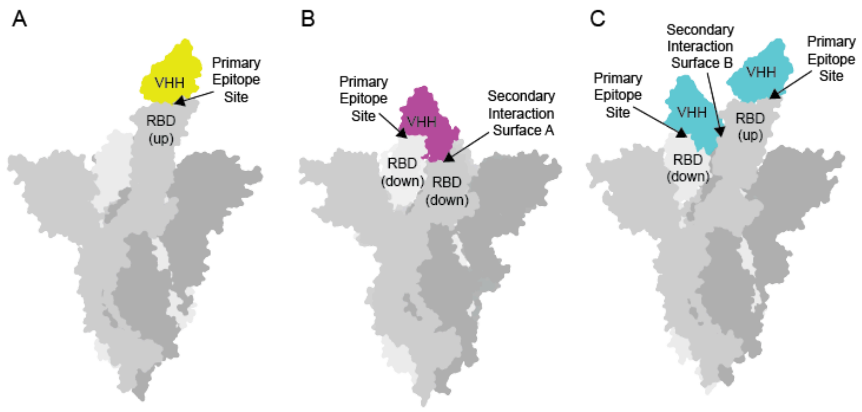


**Supplementary Figure 1. VHH binding of spike protein explained.** Three different structural classes of VHH bound to S1 trimer were determined via cryoEM. ***A***, In the class 1 structure, only one VHH binds the spike protein and recognizes an RBD in the "up" conformation. There is only one contact surface comprising the primary epitope (binding mode 1). ***B***, In the class 2 structure, as in the class 1 structure, only one VHH binds the spike protein but now recognizes the RBD in the "down" conformation. Here, there are two points of contact between the VHH and spike protein: the primary epitope site and a secondary interaction surface located on a neighboring RBD in the "down" conformation (binding mode 2). ***C***, The class 3 VHH binding scheme is a hybrid between those seen in class 1 and class 2 and has two VHH antibodies bound to the spike protein. One VHH engages an RBD in the "up" conformation through a single contact surface (binding mode 1). The other VHH recognizes an RBD in the "down" conformation and, as with class 2, there are two contact surfaces between the VHH and spike protein: the same primary epitope site on a "down" RBD and a secondary interaction surface located on a neighboring RBD but now in the "up" conformation (binding mode 3). This causes the two secondary interaction surfaces seen between class 2 and class 3 to differ slightly and involve separate sets of residues for interaction.

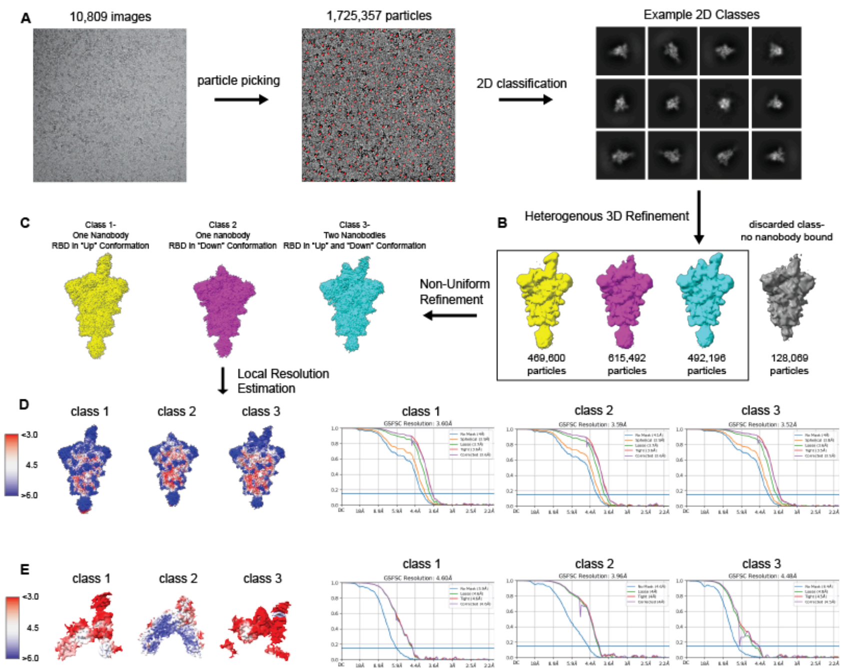


**Supplementary Figure 2. CryoEM data collection and structure Determination.** ***A***. Initial structure determination workflow. 10,809 images from data collection were subjected to particle picking where 1,725,357 particles were identified and then processed through 2D classification. These 2D classes were then subjected to heterogenous 3D refinement where 4 classes were identified. ***B***. The 3D classes produced from the initial 3D refinement revealed three classes with VHH attached to spike protein and one class showing spike only. The three VHH-containing classes were then subjected to non-uniform refinement and the non-VHH-bound class was discarded. ***C***. The resulting density maps were 3.6, 3.59, and 3.52 Å resolution. ***D***. The local resolution maps of each class showing areas of density between 3.0 and 6.0 Å and their associated FSC curves. ***E***. The local resolution maps of each class's local refinement, showing areas of density between 3.0 and 6.0 Å and their associated FSC curves. Note, in class 3, the occupancy of the VHH on the "up"-RBD is low and does not appear at a lower density threshold. Not shown, a final, composite map was created from the two density maps for each class of *D* and *E*.

**Supplementary Table 1. Cryo-EM data collection, refinement and model validation for the class 1 VHH-spike complex.**

| | | | |
|---|---|---|---|
| **Deposition** | | | |
| Coordinates (RCSB PDB) | | | |
| Map (EMDB) | | | |
| **Data Collection/ Processing** | | | |
| Microscope | TFS Titan Krios | | |
| Camera | Gatan K3 | | |
| Voltage, kV | 300 | | |
| Magnification | 81,000x | | |
| Electron exposure, $e^-/Å^2$ | 63.81 | | |
| Defocus range, µm | -0.6 to -2.6 | | |
| Pixel size, Å | 1.0691 | | |
| Symmetry | C1 | | |
| No. particles | | | |
| Initial | 1,725,357 | **Other Refinements** | |
| Final | 615,492 | **Overall** EMD-28913 | **Local** EMD-28914 |
| Map resolution (FSC 0.5), Å | 3.75 | 3.59 | 4.53 |
| **Refinement and Model** | | | |
| $CC_{mask}$ | 0.67 | | |
| Map sharpening, $Å^2$ | 83.1 | | |
| Non-hydrogen atoms | | | |
| Protein residues | 3,342 | | |
| Ligand/ modified nucleotide | 34 | | |
| B factors (min/max/mean), $Å^2$ | | | |
| Protein | 19.16/ 170.40/ 92.19 | | |
| Ligand/ modified nucleotide | 45.90/ 194.49/ 106.81 | | |
| RMS deviations | | | |
| Bond lengths, Å | 0.004 | | |
| Bond angles, ° | 0.592 | | |
| **Validation** | | | |

| | |
|---|---|
| MolProbity score | 3.09 |
| Clashscore | 9.68 |
| Poor rotamers, % | 1.21 |
| Ramachandran plot | |
| Favored, % | 93.79% |
| Allowed, % | 6.21% |
| Disallowed, % | 0% |

**Supplementary Table 2. Cryo-EM data collection, refinement and model validation for the class 2 VHH-spike complex.**

| | | Other Refinements | |
|---|---|---|---|
| **Deposition** | | | |
| Coordinates (RCSB PDB) | | | |
| Map (EMDB) | | | |
| **Data Collection/ Processing** | | | |
| Microscope | TFS Titan Krios | | |
| Camera | Gatan K3 | | |
| Voltage, kV | 300 | | |
| Magnification | 81,000x | | |
| Electron exposure, $e^-/Å^2$ | 63.81 | | |
| Defocus range, µm | -0.6 to -2.6 | | |
| Pixel size, Å | 1.0691 | | |
| Symmetry | C1 | | |
| No. particles | | | |
| Initial | 1,725,357 | **Other Refinements** | |
| Final | 489,600 | **Overall** EMD-28930 | **Local** EMD-28931 |
| Map resolution (FSC 0.5), Å | | 3.52 | 4.48 |
| **Refinement and Model** | | | |
| Model resolution (FSC 0.5), Å | 3.73 | | |
| $CC_{mask}$ | 0.76 | | |
| Map sharpening, $Å^2$ | 83.1 | | |
| Non-hydrogen atoms | | | |
| Protein residues | 3,342 | | |
| Ligand/ modified nucleotide | 37 | | |
| B factors (min/max/mean), $Å^2$ | | | |
| Protein | 15.88/ 180.54/ 87.04 | | |
| Ligand/ modified nucleotide | 37.25/ 159.76/ 92.37 | | |
| RMS deviations | | | |
| Bond lengths, Å | 0.003 | | |
| Bond angles, ° | 0.563 | | |

| | |
|---|---|
| **Validation** | |
| MolProbity score | 1.97 |
| Clashscore | 10.57 |
| Poor rotamers, % | 0.38 |
| Ramachandran plot | |
| Favored, % | 93.41% |
| Allowed, % | 6.50% |
| Disallowed, % | 0.09% |

**Supplementary Table 3. Cryo-EM data collection, refinement and model validation for the class 3 VHH-spike complex.**

| | | | |
|---|---|---|---|
| **Deposition** | | | |
| Coordinates (RCSB PDB) | | | |
| Map (EMDB) | | | |
| **Data Collection/ Processing** | | | |
| Microscope | TFS Titan Krios | | |
| Camera | Gatan K3 | | |
| Voltage, kV | 300 | | |
| Magnification | 81,000x | | |
| Electron exposure, $e^-/Å^2$ | 63.81 | | |
| Defocus range, µm | -0.6 to -2.6 | | |
| Pixel size, Å | 1.0691 | | |
| Symmetry | C1 | | |
| No. particles | | | |
| Initial | 1,725,357 | **Other Refinements** | |
| Final | 492,196 | **Overall** EMD-28868 | **Local** EMD-28880 |
| Map resolution (FSC 0.5), Å | 3.73 | 3.58 | 4.39 |
| **Refinement and Model** | | | |
| Model resolution (FSC 0.5), Å | 3.73 | | |
| $CC_{mask}$ | 0.73 | | |
| Map sharpening, $Å^2$ | 83.1 | | |
| Non-hydrogen atoms | | | |
| Protein residues | 3,460 | | |
| Ligand/ modified nucleotide | 37 | | |
| B factors (min/max/mean), $Å^2$ | | | |
| Protein | 0.09/ 485.71/ 99.29 | | |
| Ligand/ modified nucleotide | 14.83 / 171.20/ 103.88 | | |
| RMS deviations | | | |
| Bond lengths, Å | 0.003 | | |

| | |
|---|---|
| Bond angles, ° | 0.593 |
| **Validation** | |
| MolProbity score | 2.13 |
| Clashscore | 12.46 |
| Poor rotamers, % | 0.63 |
| Ramachandran plot | |
| Favored, % | 90.88% |
| Allowed, % | 8.56% |
| Disallowed, % | 0.56% |
